# Cisplatin exposure alters long-term metabolic phenotype of male, but not female, high-fat diet-fed mice

**DOI:** 10.64898/2026.04.10.717733

**Authors:** Lahari Basu, Jana Palaniyandi, Ma. Enrica Angela Ching, Myriam P. Hoyeck, Erin van Zyl, Jennifer E. Bruin

## Abstract

Cancer survivors face an increased risk of metabolic complications compared to the general population. Our group demonstrated that cisplatin, a platinum-based chemotherapeutic agent, robustly disrupts insulin secretion *in vitro* in mouse and human islets, and reduces plasma insulin levels in mice 2 weeks post-*in vivo* exposure. The long-term effects of *in vivo* cisplatin exposure alongside a pre-existing metabolic stressor, such as high-fat diet (HFD) feeding, have not been characterized. In the present study, male and female mice fed either a standard rodent chow or a 45 kcal% HFD were exposed to vehicle or 2 mg/kg cisplatin every other day for 2 weeks and then tracked for 18 weeks. Cisplatin exposure substantially influenced the metabolic phenotype of HFD-fed males but had limited impact on female HFD-fed mice. Vehicle-HFD and cisplatin-HFD male mice were both glucose intolerant compared to chow-fed controls yet, cisplatin-HFD male mice were lean, lacked a compensatory hyperinsulinemia response, and displayed increased insulin sensitivity compared to vehicle-HFD and vehicle-chow male controls. Additionally, transcriptional changes in islets isolated at 18-weeks post-exposure were largely cisplatin-driven in male mice, but diet-driven in female mice. Our study demonstrates that HFD-fed male mice exposed to cisplatin display persistent and exacerbated metabolic dysregulation relative to controls.

**ARTICLE HIGHLIGHTS:** *Why did we undertake this study?:* We previously characterized the short-term metabolic effects of cisplatin exposure *in vivo*, but the long-term metabolic effects of cisplatin remained unknown.

*What is the specific question(s) we wanted to answer?:* How does cisplatin treatment impact long-term metabolic health outcomes in mice and do outcomes differ in the presence of a metabolic stressor?

*What did we find?:* Cisplatin significantly alters the metabolic phenotype of high-fat diet-fed male mice.

*What are the implications of our findings?:* Understanding how cisplatin exposure and metabolic stress interact is critical to mitigate long-term metabolic dysregulation in cancer survivors.

## INTRODUCTION

Diabetes incidence is rising rapidly, with the International Diabetes Federation reporting an estimated 588 million individuals living with this condition (1). High-fat diet (HFD) consumption and excess body weight are well-recognized risk factors for type 2 diabetes (T2D), with HFD often leading to hyperglycemia, hyperinsulinemia, and insulin resistance (2,3). Chronic metabolic stress can lead to β-cell dysfunction, which impairs metabolic resilience and increases susceptibility to other stressors (4). Cancer survivors, regardless of age, treatment, and BMI, face an even higher risk of developing new-onset T2D compared to the general population (5–7). Interestingly, individuals who receive cisplatin—a platinum-based chemotherapeutic agent (8)—are at increased risk of developing metabolic syndrome and diabetes (7,9,10); however, the mechanisms driving this association are not fully understood.

We previously reported that cisplatin exposure reduced plasma insulin levels in male and female mice *in vivo* and potently dysregulated insulin secretion in isolated islets *ex vivo*. Cisplatin-exposed mouse islets showed supressed insulin secretion following high glucose stimulation, and heightened β-cell exocytotic capacity under low glucose conditions (11). We also observed robust downregulation of key genes linked to insulin secretion in cisplatin-exposed mouse islets compared to vehicle-exposed islets (11). Moreover, the functional and transcriptional effects of cisplatin on mouse islets were replicated in a small cohort of human donor islets (11). Although we show robust evidence of cisplatin-induced islet dysfunction, the long-term effects of cisplatin exposure *in vivo* remain unclear. Additionally, obesity and cancer treatment are both associated with secondary metabolic complications that contribute to adverse health outcomes. Thus, it is imperative to understand the effects of cisplatin on metabolic health in the presence of pre-existing metabolic stress, such as HFD. We hypothesized that HFD-feeding would exacerbate cisplatin-induced metabolic dysregulation in mice.

The goal of this study was to investigate whether pre-existing metabolic stress modifies the metabolic consequences of cisplatin exposure. To assess this, male and female mice were transferred to either fed a standard rodent chow diet or a 45 kcal% HFD for the study duration. After 4 weeks, mice were exposed to saline or cisplatin for 2 weeks (11), then monitored for 18 weeks to evaluate the long-term metabolic effects of cisplatin treatment under chow versus HFD conditions. We find that the adverse metabolic effects of cisplatin are robust and persistent in HFD-fed male mice while the effects in HFD-fed females are transient and modest. Together, our data suggest that cisplatin disrupts canonical metabolic adaptations to HFD-feeding in males but has minimal long-lasting effects in female mice.

## METHODS

### *In vivo* cisplatin exposure protocol

All experiments were approved by the Carleton University Animal Care Committee and carried out in accordance with Canadian Council on Animal Care guidelines. Male and female C57Bl/6 mice on a mixed J/N background were maintained on a 12-hour light/dark cycle with *ad libitum* access to standard rodent chow diet (Harlan Laboratories, Teklad Diet #2018) and water until ∼14 weeks of age. Experimental groups were matched for body weight and fasting blood glucose levels prior to diet switches and chemical exposure.

At ∼14 weeks of age, half of the mice of each sex were switched to a 45 kcal% HFD (Research Diets #D12451; Figure 1A) while the remaining mice continued to be fed standard chow diet. Mice were tracked on their respective diets until ∼18 weeks of age (week -2, Figure 1A), at which point they received 2 mg/kg cisplatin (Sigma-Aldrich, #232120-50MG) or 0.9% saline (vehicle control) every other day for 14 days (7 injections total; week -2 to 0, Figure 1A), as described (11). This dosing regime results in a cumulative dose of 14 mg/kg, equivalent to a 50 mg/m^2^ cisplatin administered to human cancer patients (12,13). Treatment groups included: vehicle-chow, cisplatin-chow, vehicle-HFD, and cisplatin-HFD (n=10-12/group/sex). Beginning 1-week after the cisplatin exposure period, *in vivo* metabolic assessments were conducted monthly. At 18 weeks post-exposure (∼38 weeks of age, Figure 1A), mice were euthanized for either islet isolation or tissue collection, as described below.

**Figure 1.**
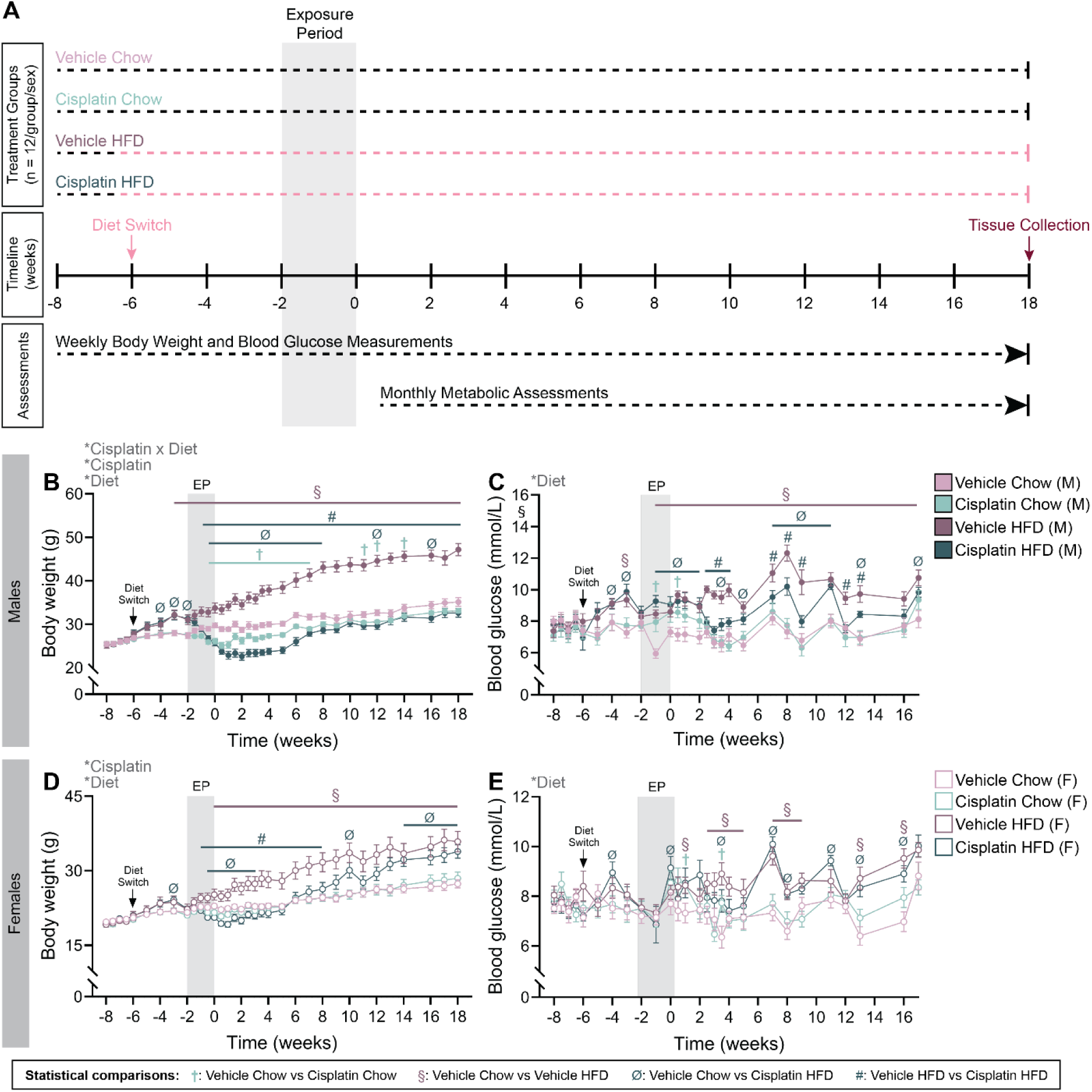
Cisplatin exposure caused weight loss and altered fasting glycemia in mice, especially in high-fat diet-fed males. **(A)** Schematic summary of the study timeline. Male and female mice were tracked for 2 weeks on a standard chow diet (week -8 to -6) after which half the mice were switched onto a 45 kcal% high-fat diet (HFD). 4 weeks post-diet switch, mice were exposed to saline (vehicle) or 2mg/kg cisplatin (n=10-12/group/sex) between weeks -2 to 0. Mice were tracked for 18 weeks post-exposure then euthanized for tissue collection. **(B, D)** Body weight and **(C, E)** fasting blood glucose in **(B-C)** male and **(D-E)** female mice. All data represented as mean ± SEM. Statistical symbols are as follows: *p < 0.05 (ANOVA main effects); ^†^p < 0.05 (vehicle chow vs. cisplatin chow); ^§^p <0.05 (vehicle chow vs vehicle HFD); ^Ø^p < 0.05 (vehicle chow vs. cisplatin HFD); ^#^p < 0.05 (vehicle HFD vs. cisplatin HFD). All graphs were analyzed using a repeated-measures 3-way mixed effects analysis with Fisher’s LSD test. M: Male, F: Female.

### *In vivo* metabolic assessments

All metabolic assessments were performed in conscious, restrained mice, and all blood sampling was conducted via saphenous vein, as previously described (14). Briefly, body weight and blood glucose levels were measured 1-2 times per week following a 4-hour morning fast. Blood glucose was measured using the MediSure Multi-Patient Use Glucose Meter (MediSure Canada, measuring range: 1.1–41.7 mmol/L) from study weeks -8 to 2, and using the StatStrip XPRESS Glucose Meter (Nova Biomedical, measuring range: 0.5–33.3 mmol/L) from study week 3 onward.

For metabolic assessments, 0 minutes indicates data collected prior to administration of glucose or insulin. For ip glucose tolerance tests (ipGTTs), mice received an ip injection of glucose (1-2 g/kg; DMVet Dextrose 50%, #02420880) following a 6-hour morning fast. Plasma insulin levels were measured by ELISA (ALPCO, #80-INSMSU-E10). For insulin tolerance tests (ITTs), mice received an ip injection of insulin (0.6-0.7 IU/kg; Novolin ge Toronto, Novo Nordisk, #02024233) following a 4-hour morning fast. For ipGTTs and ITTs, the MediSure Multi-Patient Use Glucose Meter was used on weeks 1-2 and the OneTouch Verio Flex (LifeScan, measuring range: 1.1-33.3 mmol/L) was used from week 6 onward.

### Mouse islet isolation and tissue collection

Islets were isolated from a subset of mice (n=6-7/group/sex) and handpicked to >99% purity, as previously described (11). Islets were incubated overnight in RPMI 1640 media (Wisent Bioproducts, #350-000-CL) supplemented with 10% (vol./vol.) FBS (Sigma-Aldrich, #F1051-500ML) and 1% (vol./vol.) penicillin-streptomycin (Gibco, #15140-122-100) at 37°C with 5% CO_2_ prior to functional analyses. From the remaining mice (n=4-5/group/sex), intact pancreas was collected for electron microscopy and paraffin-embedding, and liver was flash frozen and collected for RNA extraction (described below).

### *Ex vivo* glucose-stimulated insulin secretion assays

To assess static glucose-stimulated insulin secretion in isolated islets, 25 islets/replicate (n=3 technical replicates/mouse, n=4-5 mice/group/sex) were handpicked and underwent sequential 1-hour incubations in Krebs-ringer bicarbonate HEPES buffer (KRBH) containing 2.8 mmol/L glucose (low glucose; LG), 16.7 mmol/L glucose (high glucose, HG), and 30 mmol/L KCl KRBH, as previously described (14).

### Oxygen consumption analysis

To study mitochondrial function and islet respiration, 70 islets/mouse (n=4-5 mice/group/sex) were handpicked, incubated in Seahorse XF RPMI media (Agilent Technologies, 103576, pH 7.4) supplemented with 2 mmol/L sodium pyruvate, 2 mmol/L L-glutamine, and 1% (vol./vol.) FBS, and 2.8 mmol/L glucose, and used to measure oxygen consumption rate using a Seahorse XFe24 Analyzer (Agilent Technologies) as previously described (11).

### Transmission electron microscopy (TEM)

Intact pancreatic tissue was cut into small, ∼1mm long pieces and stored in 2% (v/v) glutaraldehyde (Sigma, #G5882-10X1ML) in 0.2M sodium cacodylate buffer (#CA100503-718) at 4°C. Fixed pancreas samples were processed and sectioned by the Electron Microscopy Facility at McMaster University (Hamilton, Ontario, Canada). Tissues were transferred to 2% (v/v) glutaraldehyde (#16120) in 0.1M sodium cacodylate (#12300), washed twice with buffer solution, then fixed in 1% OsO4 in 0.1M sodium cacodylate buffer for 1 hour at room temperature. Tissues were then dehydrated in an ethanol series (50%, 70% twice, 95% twice, 100% twice), followed by 2 washes in 100% propylene oxide (PO, #20411), and gradual infiltration through a graded Spurr’s resin (#14300) series (2:1 PO:Spurr’s, 1:1 PO:Spurr’s, 1:2 PO:Spurr’s, 100% Spurr’s three times) while rotating samples. Infiltrated samples were transferred to embedding moulds filled with fresh 100% Spurr’s and polymerized overnight at 60°C. Thin sections (80-90 nm) were cut using an Ultracut UCT ultramicrotome (Leica Microsystems) and placed onto 200-mesh Cu/Pd grids (#G200-CP). Sections were stained with UranyLess (#22409) and lead citrate (#22410). All supplies used for TEM were purchased from Electron Microscopy Sciences, unless otherwise stated. Grids were examined and imaged with an FEI Tecnai G2 F20 TEM (FEI Co.). β-cells were identified within the pancreas sections by the presence of dense-core insulin granules.

### Histology

Intact pancreas tissue was collected and fixed in 4% paraformaldehyde (PFA; Fisher Scientific, #AAJ19943K2) for 24 hours, then stored in 70% (v/v) ethanol. PFA-fixed pancreas tissues were processed and paraffin-embedded by the University of Ottawa Heart Institute Histology Core Facility (Ottawa, Ontario, Canada). Immunofluorescent staining was performed as described (15). Primary and secondary antibodies are listed in Supplementary Table 1.

Pancreas sections were imaged using an Axio Observer 7 microscope (Carl Zeiss) and quantified using Zen Blue 2.6 software (Carl Zeiss). The percentage of hormone^+^ area was calculated as (average [hormone^+^ area/ islet^+^ area of each islet in a biological replicate] x 100%). Only islets ≥5000 μm^2^ were included in this analysis. The percentage of β-cells with cytoplasmic proinsulin accumulation was calculated as ([total insulin^+^ cells with cytoplasmic proinsulin accumulation/total insulin^+^ cells in a biological replicate] x 100%). Similarly, the percentage of insulin^+^MAFA^+^ cells was calculated as ([total insulin^+^MAFA^+^ cells/total insulin^+^ cells in a biological replicate] x 100%). For these measurements, only islets with >30 cells and >400 total insulin^+^ cells were included.

### Quantitative real time PCR

To evaluate changes in mRNA expression, RNA was isolated from mouse islets (90-381 islets/mouse) stored in buffer RLT supplemented with 1% β-mercaptoethanol, using the RNeasy Micro Kit (Qiagen, #74004) following the manufacturer’s protocol. For liver samples, RNA was extracted from flash-frozen tissue using TRIzol™ reagent (Invitrogen, #15596018), as per manufacturer’s instructions. DNase treatment was performed prior to cDNA synthesis using the iScript gDNA Clear cDNA synthesis Kit (Bio-Rad, #1725035). qPCR was performed using SsoAdvanced Universal SYBR Green Supermix (Bio-Rad, #1725271) and run on a CFX384 (Bio-Rad). All targets were quantified alongside “no reverse transcriptase” and “no cDNA template” controls. *Ppia* was used as the reference gene for islets, while the average Ct values of *Ppia* and *Gapdh* were used as the housekeeping reference for liver. Data were analyzed using the 2^-ΔΔCt^ method. Primer sequences are listed in Supplementary Table 2.

### Statistical Analysis

All statistical analyses were conducted using GraphPad Prism 10.6.0 (GraphPad Software). Specific statistical tests and sample sizes are indicated in figure legends. For all analyses, p<0.05 was considered statistically significant. Data are presented as mean ± SEM.

### Data and Resource Availability

All data that support the findings of this study are available from the corresponding author upon reasonable request.

## RESULTS

### Cisplatin exposure reduced body weight and altered fasting blood glucose levels in mice

Cisplatin-exposed male mice experienced significant weight loss compared to vehicle-exposed controls, but the magnitude and persistence of weight loss varied based on diet (Figure 1B). Cisplatin-chow male mice lost ∼8% of body weight during the exposure period while cisplatin-HFD males lost ∼15% body weight (Figure 1B). Cisplatin-chow male mice regained weight and were comparable to vehicle-chow mice by the end of the study (Figure 1B). In contrast, cisplatin-HFD males did not gain weight at a similar rate to vehicle-HFD males and by the end of the study, were ∼30% lighter than vehicle-HFD males and similar weight to chow-fed males (Figure 1B). Male cisplatin-chow mice had transiently elevated fasting blood glucose compared to vehicle-chow males at 1 week into the exposure period (week -1) and at week 0.5 (Figure 1C). Interestingly, cisplatin-HFD males had sporadic patterns in their fasting glycemia, and only consistently matched vehicle-HFD blood glucose levels 16 weeks after the exposure period (Figure 1C).

Cisplatin exposure did not affect body weight in chow-fed female mice. Cisplatin-HFD female mice weighed ∼ 15% less than vehicle-HFD females between weeks -1 (i.e., mid-exposure period) and 8, but regained body weight thereafter and were comparable to vehicle-HFD females by week 9 (Figure 1D). Cisplatin-chow female mice had modestly elevated fasting blood glucose levels compared to vehicle-chow females at weeks 1 and 3.5, but HFD-feeding, irrespective of chemical, drove intermittent fasting hyperglycemia in female mice compared to chow-fed females between weeks 7 to 17 (Figure 1E).

### Cisplatin-exposed male mice lacked a compensatory hyperinsulinemia response to HFD-feeding and were increasingly insulin sensitive

Cisplatin-chow males were transiently glucose intolerant relative to vehicle-chow controls at weeks 1 and 10 post-exposure (Figure 2A-B), but their glucose tolerance normalized by week 18 (Figure 2C). In contrast, cisplatin-HFD males had transiently decreased blood glucose during a GTT at week 10 compared to vehicle-HFD males (Figure 2B). Importantly, HFD-fed males from both treatment groups were glucose intolerant compared to chow-fed controls starting as early as 1-week post-exposure (Figure 2A-C).

**Figure 2.**
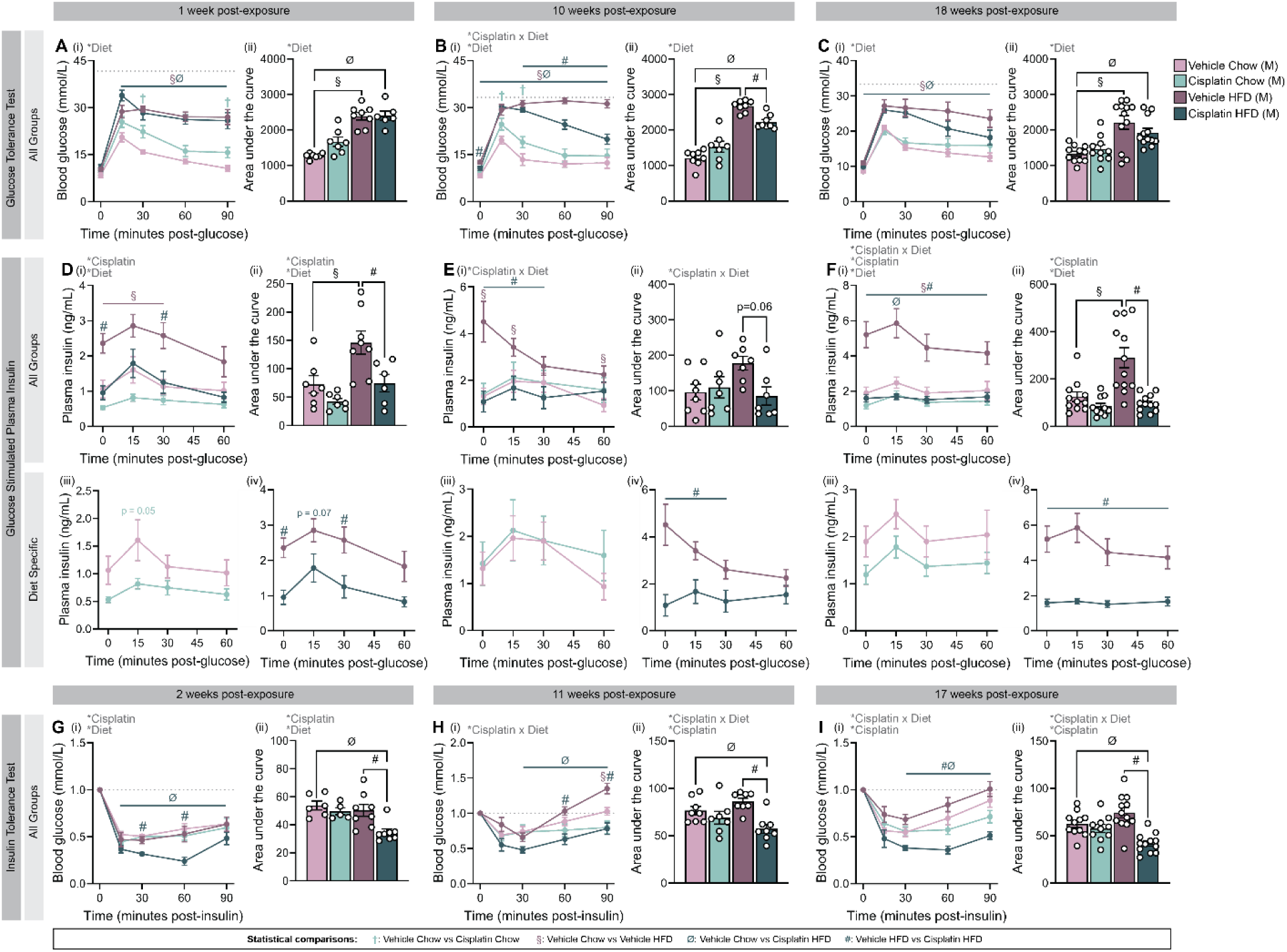
Cisplatin-HFD male mice show lower plasma insulin levels and increased insulin sensitivity compared to vehicle-HFD males, but remain glucose intolerant relative to chow-fed controls. (A-C) Glucose tolerance and **(D-F)** glucose-stimulated plasma insulin levels were measured in male mice at **(A, D)** 1, **(B, E)** 10, and **(C, F)** 18 weeks post-exposure; mice received intraperitoneal (ip) injections of **(A, D)** 2 g/kg glucose, or **(B, C, E, F)** 1 g/kg glucose. **(G-I)** Insulin tolerance tests were conducted at **(G)** 2, **(H)** 11, and **(I)** 17 weeks post-exposure; blood glucose values are normalized to baseline (t=0). Mice received ip injections of **(G)** 0.7 IU/kg insulin, or **(H-I)** 0.6 IU/kg insulin. **(A-C)** Glucose tolerance and **(G-I)** insulin tolerance are presented as (i) line graphs and (ii) area under the curve. **(D-F)** Plasma insulin levels presented as (i) line graphs with all groups, (ii) area under the curve, (iii) line graphs with only chow-fed groups, and (iv) line graphs with only HFD-fed groups. All data represented as mean ± SEM. Statistical symbols are as follows: *p < 0.05 (ANOVA main effects); ^†^p < 0.05 (vehicle chow vs. cisplatin chow); ^§^p <0.05 (vehicle chow vs vehicle HFD); ^Ø^p < 0.05 (vehicle chow vs. cisplatin HFD); ^#^p < 0.05 (vehicle HFD vs. cisplatin HFD). The following statistical tests were used: line graphs (A, F) repeated measures mixed-effects analysis with Fisher’s LSD test; (B-E, G-I) repeated measures 3-way ANOVA with Fisher’s LSD test; all bar graphs analyzed with ordinary 2-way ANOVA with Tukey’s multiple comparisons test. M: Male.

Both cisplatin-chow and cisplatin-HFD males demonstrated reduced basal and glucose-stimulated plasma insulin levels compared to their respective diet controls at 1-week post-exposure (Figure 2D). Cisplatin-chow males restored plasma insulin levels to match vehicle-chow mice over time (Figures 2Eiii, Fiii, and S1Ciii, Diii). In contrast, cisplatin-HFD mice continued to have a ∼2-fold reduction in plasma insulin levels compared to vehicle-HFD mice throughout the study (Figures 2Eiv, Fiv, and S1Civ, Div). Ultimately, cisplatin-HFD males failed to mount compensatory hyperinsulinemia for the duration of the study despite their continued glucose intolerance (Figure 2D-F, S1C-D).

Surprisingly, cisplatin-HFD male mice had increased insulin sensitivity compared to both vehicle-HFD and vehicle-chow males at weeks 2, 7, and 11 (Figures 2G-I and S1E), despite persistent glucose intolerance (Figures 2A-C and S1A-B). In fact, cisplatin-HFD males continued to become progressively more insulin sensitive, culminating to a 2-fold decrease in blood glucose levels post-insulin compared to vehicle-HFD males at week 17 (Figure 2Iii).

To better understand the direct effects of cisplatin on islet function *in vivo*, islets isolated from male mice 18 weeks post-exposure were used to assess glucose-stimulated insulin secretion (GSIS) and oxygen consumption *ex vivo* (Figure S3A-K). Contrary to the robust changes in plasma insulin levels *in vivo* (Figure 2D-F), we did not observe any changes in GSIS, stimulation index, or insulin content of the islets between treatment groups (Figure S3A-C). No profound changes in oxygen consumption were observed (Figure S3D-K), but there was a modest overall effect of cisplatin to decrease spare respiratory capacity of islets from male mice (Figure S3H).

Overall, cisplatin induced metabolic dysregulation in male mice, though its effects varied based on diet. While cisplatin-chow male mice recovered relatively quickly from their initial hyperglycemia and decreased plasma insulin levels, cisplatin-HFD mice exhibited prolonged metabolic dysregulation (Figure 2, S1). Paradoxically, cisplatin-HFD mice were markedly more insulin sensitive than all other groups (Figure 2G-I, S1E), suggesting a dissociation between glucose tolerance and insulin action.

### Cisplatin exposure modestly, and acutely, alters metabolic function in female mice

The effects of cisplatin on metabolic homeostasis were more modest in female mice. Interestingly, at week 18, cisplatin-chow females exhibited mildly elevated fasting blood glucose compared to vehicle-chow females (Figure 3C) but did not demonstrate altered glycemia at any prior timepoint. Overall, cisplatin-exposed females had comparable glucose tolerance to their respective diet controls (Figures 3A-C, S2A-B). In line with the GTT data, cisplatin exposure had no effect on glucose-stimulated plasma insulin levels throughout the study (Figures 3D, F, and S2C, D), aside from trending increase in insulin at 15-minutes post-glucose at week 10 in cisplatin-chow females (Figure 3E). In contrast, cisplatin-HFD females displayed an overall reduction in glucose-stimulated plasma insulin levels compared to vehicle-HFD females at week 1 (Figure 3Dii and 3Div), but this effect was transient and not observed at week 10 or 18 (Figure 3E-F). Female cisplatin-exposed mice also showed modest and transient differences in insulin sensitivity throughout the study. Cisplatin-chow females were transiently more insulin sensitive than vehicle-chow females at weeks 7 and 10, but this effect was very modest (Figure 2Hi, S2Ei). Cisplatin-HFD females had modestly decreased blood glucose levels at isolated timepoints during ITTs at 2 and 11 weeks post-exposure (Figures 3G and 3H), and a robust increase in insulin sensitivity 7 weeks post exposure (Figure S2E), but no differences at week 17 (Figure 3I) compared to vehicle controls.

**Figure 3.**
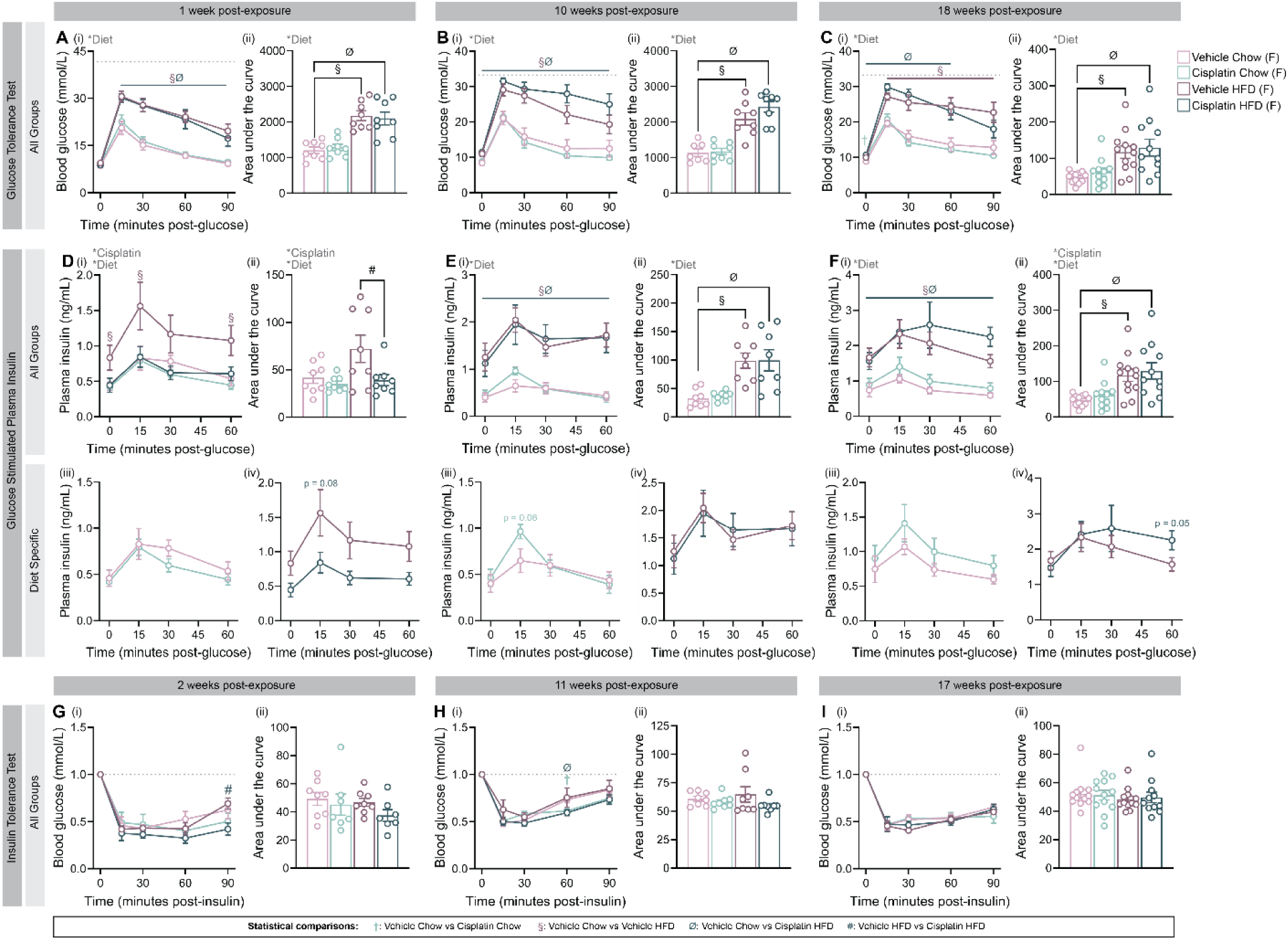
Glucose homeostasis was primarily affected by diet, not cisplatin, in female mice. (A-C) Glucose tolerance and **(D-F)** glucose-stimulated plasma insulin levels were measured in female mice at **(A, D)** 1, **(B, E)** 10, and **(C, F)** 18 weeks post-exposure; mice received intraperitoneal (ip) injections of **(A, D)** 2 g/kg glucose, or **(B, C, E, F)** 1 g/kg glucose. **(G-I)** Insulin tolerance tests were conducted at **(G)** 2, **(H)** 11, and **(I)** 17 weeks post-exposure; blood glucose values are normalized to baseline (t=0). Mice received ip injections of **(G)** 0.7 IU/kg insulin, or **(H-I)** 0.6 IU/kg insulin. **(A-C)** Glucose tolerance and **(G-I)** insulin tolerance are presented as (i) line graphs and (ii) area under the curve. **(D-F)** Plasma insulin levels presented as (i) line graphs with all groups, (ii) area under the curve, (iii) line graphs with only chow-fed groups, and (iv) line graphs with only HFD-fed groups. All data represented as mean ± SEM. Statistical symbols are as follows: *p < 0.05 (ANOVA main effects); ^†^p < 0.05 (vehicle chow vs. cisplatin chow); ^§^p <0.05 (vehicle chow vs vehicle HFD); ^Ø^p < 0.05 (vehicle chow vs. cisplatin HFD); ^#^p < 0.05 (vehicle HFD vs. cisplatin HFD). The following statistical tests were used: line graphs, repeated measures 3-way ANOVA with Fisher’s LSD test; bar graphs, 2-way ANOVA with Tukey’s multiple comparisons test. F: Female.

In line with our *in vivo* assessments, ex *vivo* analyses of isolated islets from female mice revealed no differences in GSIS, stimulation index, insulin content, or overall oxygen consumption between treatment groups (Figure S3L-O). Although pairwise comparisons revealed no statistically significant differences in oxygen consumption parameters between treatment groups (Figure S3P-V), an overall significant effect of diet led to reduced maximal respiration and mitochondrial ATP production in islets from HFD-fed females compared to chow-fed females (Figure S3Q, T).

Overall, female mice from both diet groups were largely protected from the long-term adverse metabolic effects of cisplatin, unlike males. While the changes in metabolic function of male mice were primarily driven by cisplatin, the changes observed in female mice were largely attributable to diet-induced effects.

### Cisplatin-HFD males did not increase islet size, while cisplatin-HFD females had altered proinsulin immunoreactivity

To determine if the decreased plasma insulin levels observed in cisplatin-exposed male mice were driven by protein-level changes, we assessed islet morphology and other β-cell characteristics in pancreas tissue at 18 weeks post-exposure. Cisplatin did not alter the %insulin^+^ or %glucagon^+^ area per islet in male mice (Figure 4A-B, L). Qualitative assessments of transmission electron microscopy images showed no marked differences in insulin content within β-cells of cisplatin-exposed vs vehicle-exposed male mice (Fig 4K), consistent with the normal immunofluorescent staining for insulin in paraffin-embedded pancreas (Figure 4A) and measurement of total insulin content in lysed isolated islets (Figure S3C). However, cisplatin-HFD males failed to exhibit the characteristic islet hypertrophy observed in vehicle-HFD males relative to chow-fed controls, suggesting a blunted islet adaptive response to HFD (Figure 4C).

**Figure 4.**
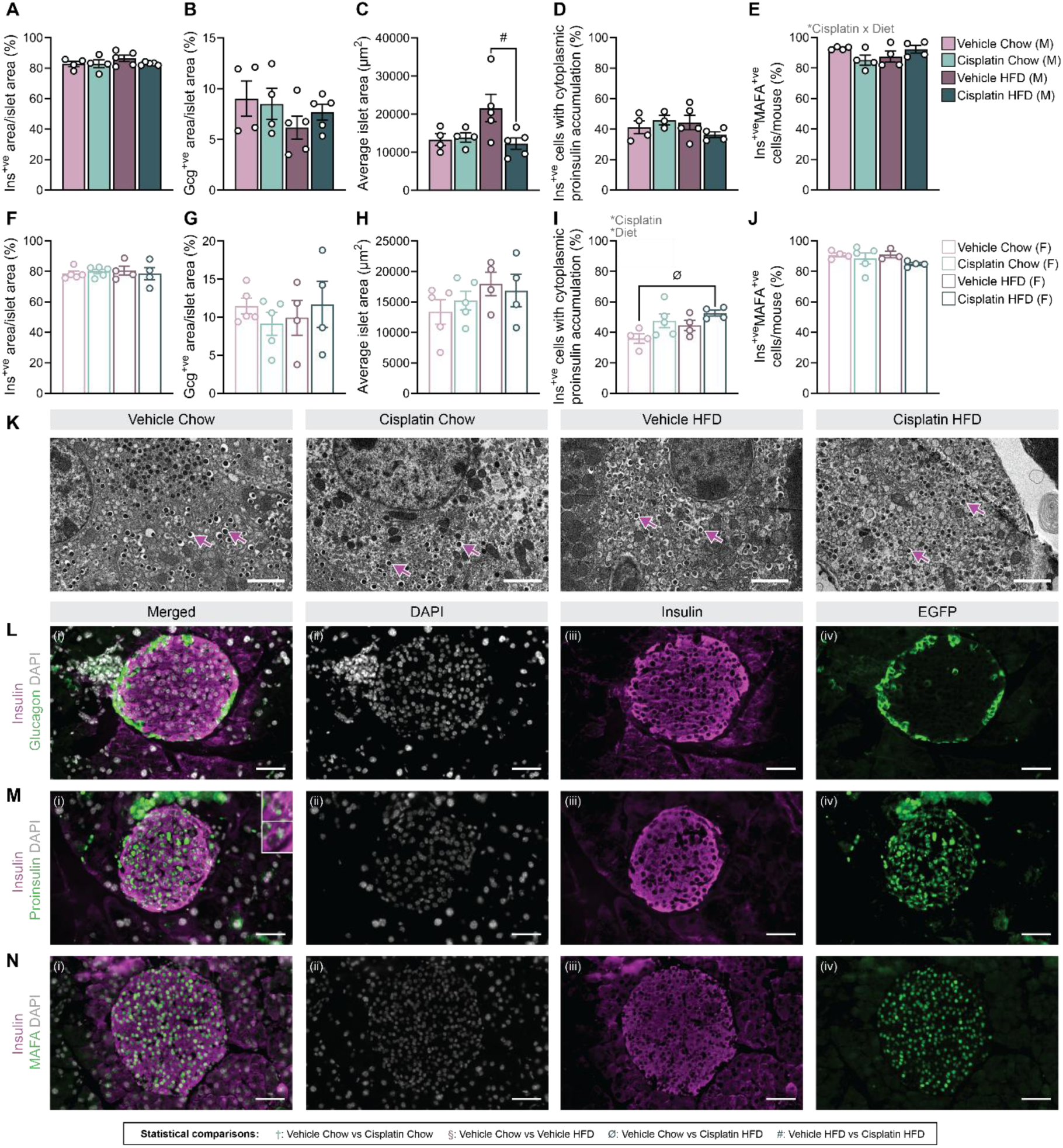
Cisplatin caused modest changes in islet composition in both male and female HFD-fed mice. Whole pancreas harvested from **(A-E)** male and **(F-J)** female mice 18 weeks post-exposure were used to assess islet morphology. **(A, F)** Percentage of insulin^+^ area per islet, **(B,G)** percentage of glucagon^+^ area per islet, **(C, H)** total islet area, **(D, I)** percentage of β-cells with cytoplasmic proinsulin accumulation, and **(E, J)** percentage of Ins^+^/MAFA^+^ cells per mouse as determined by immunofluorescent staining. **(K)** Representative electron microscopy photographs of pancreatic islets. Magenta arrows indicate insulin granules. Scale bar = 2 μm. **(L-N)** Representative images of pancreas section showing immunofluorescence staining of **(L)** insulin and glucagon, **(M)** insulin and proinsulin, and **(N)** insulin and MAFA. Inset regions of **(M)** show examples of perinuclear (top) and cytoplasmic (bottom) accumulation of proinsulin. Representative images shown as (i) all channels merged, (ii) DAPI, (iii) insulin, and (iv) EGFP. Scale bar = 50 μm. All data represented as mean ± SEM. Statistical symbols are as follows: *p < 0.05 (ANOVA main effects); ^†^p < 0.05 (vehicle chow vs. cisplatin chow); ^§^p <0.05 (vehicle chow vs vehicle HFD); ^Ø^p < 0.05 (vehicle chow vs. cisplatin HFD); ^#^p < 0.05 (vehicle HFD vs. cisplatin HFD). All graphs analyzed using a 2-way ANOVA with Tukey’s multiple comparisons test. M: Male, F: Female.

To explore if improper insulin processing contributed to reduced plasma insulin levels in cisplatin-HFD males, we quantified insulin^+^ cells with cytoplasmic proinsulin accumulation but found no differences between treatment groups (Figure 4D, M). To determine if changes in β-cell identity were a driving factor in the reduction of plasma insulin, we measured the presence of MAFA in insulin^+^ cells. There were no differences in the percentage of insulin^+^MAFA^+^ cells between treatment groups in male mice (Figure 4E, N).

In female mice, there were no changes in % insulin^+^ area, or % glucagon^+^ area, average islet area, or % insulin^+^MAFA^+^ cells (Figure 4F-H, J), but there was a significant increase in cytoplasmic proinsulin accumulation in cisplatin-HFD β-cells compared to vehicle-chow β-cells (Figure 4I). Interestingly, cisplatin contributed to a significant overall increase in cytoplasmic proinsulin accumulation relative to vehicle exposure in female mice (Figure 4I).

Our results suggest that the pronounced decrease in plasma insulin levels in cisplatin-HFD males relative to vehicle-HFD males is not driven by changes in islet composition, insulin content, or β-cell identity. However, consistent with their inability to compensate for hyperglycemia, cisplatin-HFD male mice did not increase their islet size in response to HFD-feeding, unlike vehicle-HFD controls. While there were no overt differences observed in metabolic function *in vivo* in female mice, the increased accumulation of cytoplasmic proinsulin suggests cisplatin alters insulin protein processing, which could have long-term consequences on metabolic health beyond 18 weeks post-exposure.

### Cisplatin alters expression of genes involved in insulin processing, hormone production, and β-cell identity in male, but not female, mice

We next explored the effects of cisplatin exposure and HFD-feeding on islets at the transcriptional level, including key genes involved in insulin processing, hormone production, and β-cell identity, at 18 weeks post-exposure (Figures 5, 6). *Ins1* expression was decreased ∼2-fold in islets from cisplatin-chow compared to vehicle-chow male mice, but was not different between cisplatin-HFD versus vehicle-HFD mice (Figure 5A). In contrast, cisplatin increased *Gcg* expression by ∼2-fold increase in in islets from HFD-fed, but not chow-fed, males (Figure 5D). Cisplatin also reduced the expression of both *Pcsk1* and *Pcsk2* in islets from male mice, regardless of diet (Figure 5E, F). No changes in *Ins2*, *Sst*, or *Slc30a8* (zinc transporter) expression were observed in male islets from any treatment group (Figure 5B, C, F). Together, this suggests that cisplatin may exert its effects at a transcriptional level on insulin production and processing.

**Figure 5.**
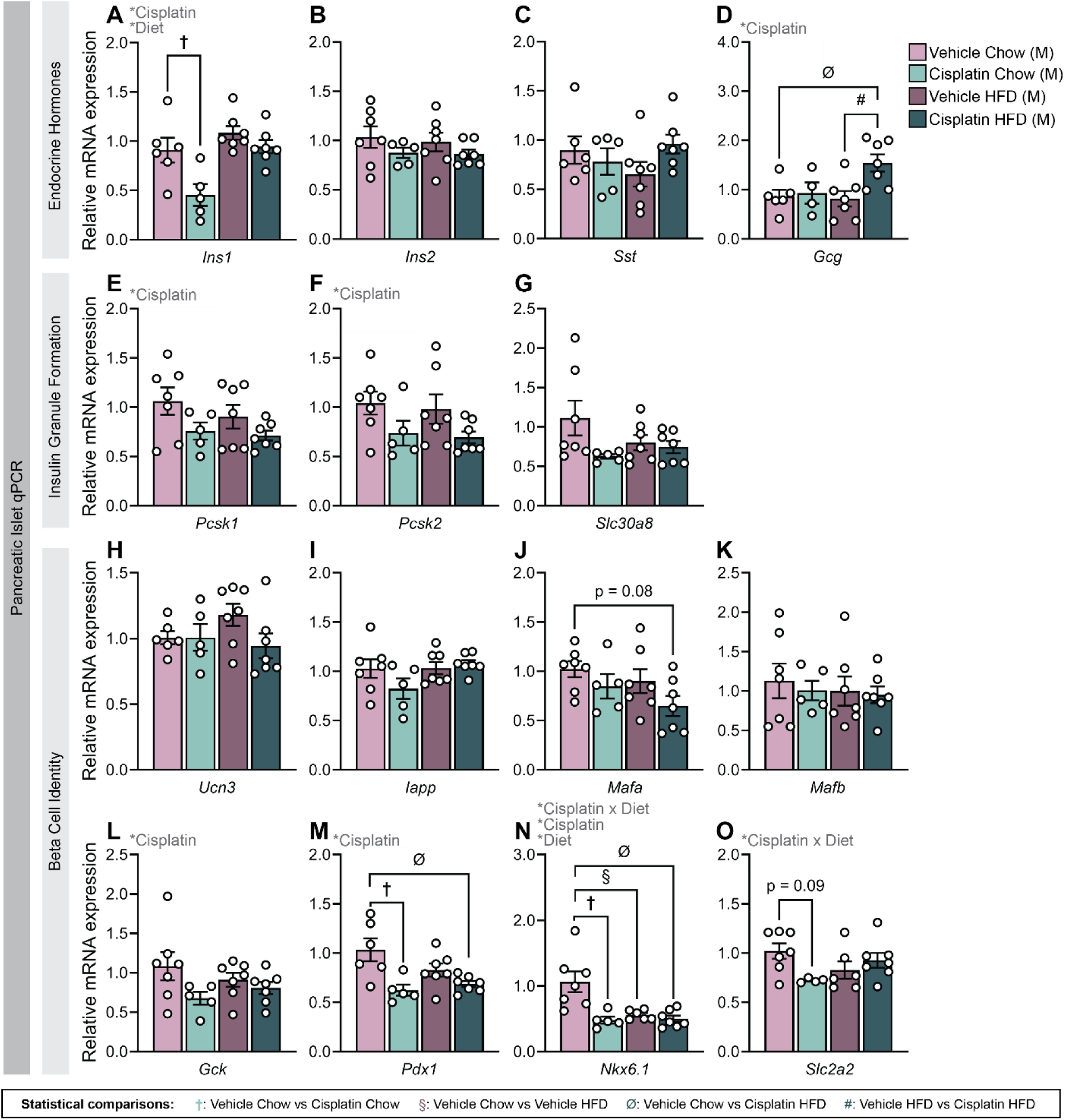
Cisplatin exposure had significant overall effects on insulin processing and beta cell identity genes in islets from male HFD-fed mice. Islets isolated from male mice 18 weeks post exposure were used to measure relative mRNA expression of (A) *Ins1*, (B) *Ins2*, (C) *Sst*, (D) *Gcg*, (E) *Pcsk1*, (F) *Pcsk2*, (G) *Slc30a8*, (H) *Ucn3*, (I) *Iapp*, (J) *Mafa*, (K) *Mafb*, (L) *Gck*, (M) *Pdx1*, (N) *Nkx6-1*, and (O) *Slc2a2*. All data represented as mean ± SEM. Statistical symbols are as follows: *p < 0.05 (ANOVA main effects); ^†^p < 0.05 (vehicle chow vs. cisplatin chow); ^§^p <0.05 (vehicle chow vs vehicle HFD); ^Ø^p < 0.05 (vehicle chow vs. cisplatin HFD); ^#^p < 0.05 (vehicle HFD vs. cisplatin HFD). All graphs analyzed using 2-way ANOVA with Tukey’s multiple comparisons test. M: Male.

We next evaluated β-cell identity and maturity markers in male islets (Figure 5H-O). Expression of *Ucn3, Iapp,* and *Mafb* were unchanged across all groups (Figure 5H, I, K). *Mafa* expression was modestly, but not significantly, reduced in cisplatin-HFD islets compared to vehicle-chow controls (Figure 5J); there were no other robust differences in *Mafa* expression, aligning with our quantification of MAFA at the protein level (Figure 4E). *Gck* expression was modestly decreased in cisplatin-exposed male islets overall compared to vehicle-exposed islets (Figure 5L). Notably, both *Pdx1* and *Nkx6-1* expression were downregulated in cisplatin-HFD islets and cisplatin-chow islets compared to vehicle-chow islets (Figure 5M, N). Finally, cisplatin-chow male islets showed a modest reduction in *Slc2a2* expression (encoding the glucose transporter GLUT2) compared to vehicle-chow islets; there was no effect of cisplatin on *Slc2a2* expression in HFD-fed males (Figure 5O).

Given the elevated *Gcg* expression in cisplatin-HFD male islets, we next asked whether altered glucagon signaling might contribute to the impaired glucose tolerance observed in these mice. We examined genes regulating gluconeogenesis and glycogenolysis in the liver (Figure S4A-D). Neither liver *Gcgr* (glucagon receptor) nor *Pck1* (key gluconeogenic enzyme) expression differed across groups (Figure S4A, B), but *G6pc* (a key gluconeogenesis enzyme) was significantly reduced in cisplatin-HFD males compared to vehicle-chow males (Figure S4C).

In female mice, gene expression changes in islets and liver were more modest (Figures 6, S4E-H). Cisplatin-HFD female islets had increased *Ins1* expression compared to vehicle-chow (Figure 6D), while cisplatin increased *Gcg* irrespective of diet (Figure 6G). Moreover, while diet caused a modest decrease in *Mafa* expression in female islets, there were no effects of cisplatin on *Mafa* or other β-cell identity markers (Figure 6H-O).

**Figure 6.**
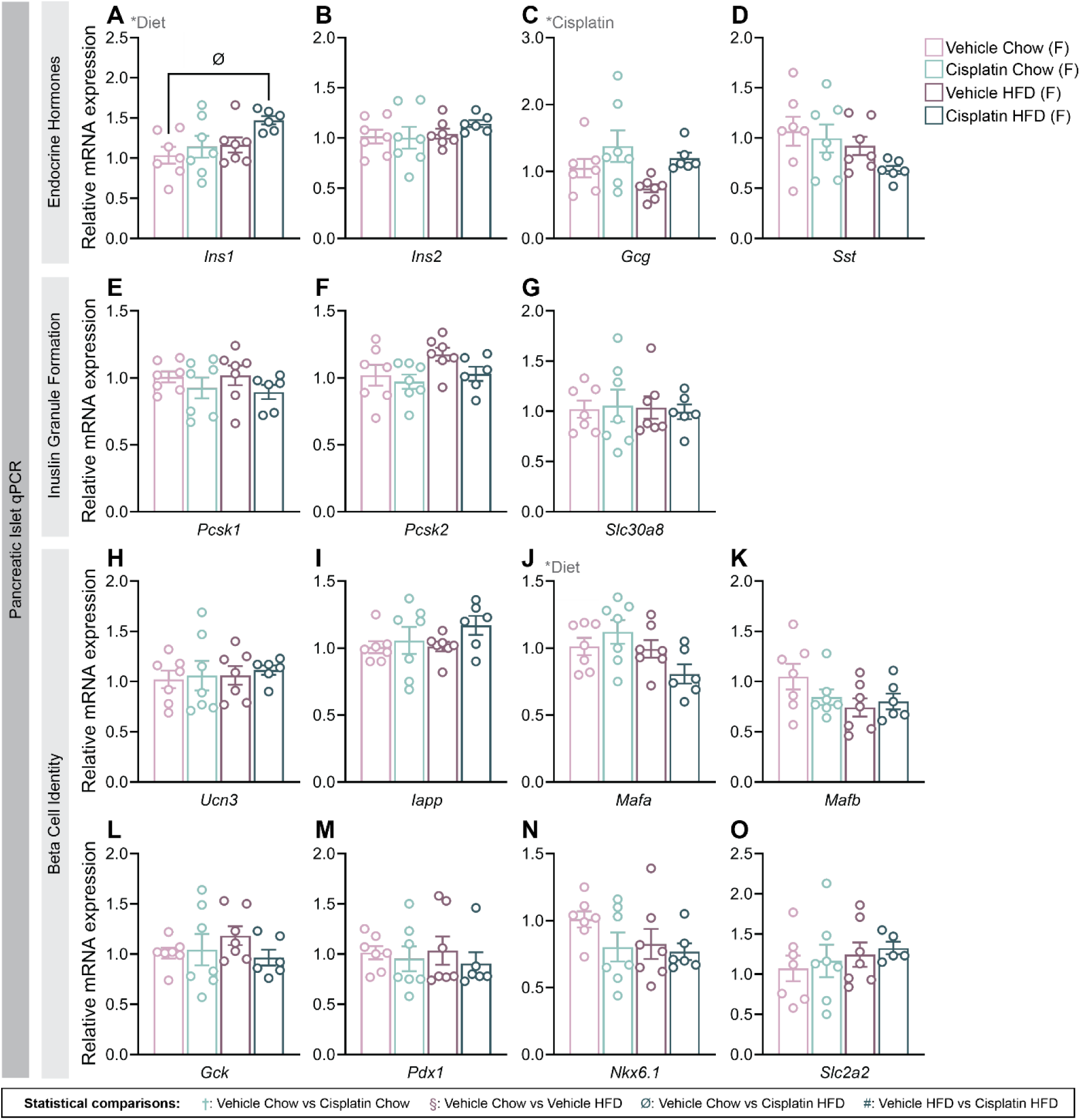
Expression of key genes linked to islet function were largely unaffected by cisplatin or diet in female mice. Islets isolated from female mice 18 weeks post exposure were used to measure relative mRNA expression of **(A)** *Ins1*, **(B)** *Ins2*, **(C)** *Sst*, **(D)** *Gcg*, **(E)** *Pcsk1*, **(F)** *Pcsk2*, **(G)** *Slc30a8*, **(H)** *Ucn3*, **(I)** *Iapp*, **(J)** *Mafa*, **(K)** *Mafb*, **(L** *Gck*, **(M)** *Pdx1*, **(N)** *Nkx6-1*, and **(O)** *Slc2a2*. All data represented as mean ± SEM. Statistical symbols are as follows: *p < 0.05 (ANOVA main effects); ^†^p < 0.05 (vehicle chow vs. cisplatin chow); ^§^p <0.05 (vehicle chow vs vehicle HFD); ^Ø^p < 0.05 (vehicle chow vs. cisplatin HFD); ^#^p < 0.05 (vehicle HFD vs. cisplatin HFD). All graphs analyzed using 2-way ANOVA with Tukey’s multiple comparisons test. F: Female.

Overall, we found that although cisplatin did not overtly alter insulin content or proinsulin accumulation in islets from male mice (Figure S3C and 4D), cisplatin broadly supressed transcription of genes essential for β-cell function and identity in male islets. In female mice, diet, rather than cisplatin exposure, was the primary driver of the modest transcriptional changes observed. Together, these findings suggest cisplatin alters both β- and α-cell gene networks, with more pronounced effects in male mice than female.

## DISCUSSION

Cancer survivors treated with cisplatin-based chemotherapy have an increased risk of metabolic dysfunction compared to the general population (7,16). Characterizing the interactions between cisplatin and diet-induced metabolic stress is imperative to better understand the metabolic consequences that can influence long-term health outcomes of cancer survivors. We show that cisplatin exposure profoundly alters metabolic adaptation to HFD-feeding in male mice, leading to long-lasting impairments of glucose tolerance, insulin secretion, and diet-induced weight gain. Conversely, the effects of cisplatin in female mice were transient, even in HFD-fed animals. Our study suggests that cisplatin disrupts the coordinated relationship between insulin secretion, insulin action, and islet plasticity that normally supports adaption to HFD-feeding in a sex-dependent manner.

The most striking, and paradoxical, finding from this study was the metabolic phenotype displayed by cisplatin-HFD males—low body weight, glucose intolerance, a lack of compensatory hyperinsulinemia, and increased insulin sensitivity. Canonically, HFD-feeding in C57Bl/6 mice leads to weight gain, glucose intolerance, and compensatory hyperinsulinemia (17), all of which were observed in vehicle-HFD male and HFD-fed female mice; in contrast, cisplatin-HFD males exhibited glucose intolerance but none of the other canonical HFD-induced metabolic phenotypes.

The unique metabolic phenotype exhibited by cisplatin-HFD male mice resembles “type 5 diabetes”, which is characterized by low BMI, severe insulin deficiency, and moderate to severe hyperglycemia (18,19). A key feature in type 5 diabetes is defective insulin secretion, rather than peripheral insulin resistance, leading to improper glycemic control (20), further aligning with the phenotype observed in cisplatin-HFD male mice.

An interesting aspect of the metabolic phenotype in male cisplatin-HFD mice was their inability to gain weight compared to vehicle-HFD mice. Although lean and fat mass were not measured in these mice, cisplatin-HFD male mice were ∼30% lighter than vehicle-HFD male mice at the end of the study. Similarly, cisplatin-HFD female mice were ∼20% lighter than vehicle-HFD females between 0-10 weeks post-exposure. Cisplatin is known to exert cytotoxic effects on the gastrointestinal tract and kidneys (21–23), which can impair proper nutrient absorption. The difference in body weight could suggest the possibility of impaired lipid storage capacity and altered nutrient handling by cisplatin-exposed mice (24,25). Interestingly, Mehran *et al.* suggested that hyperinsulinemia is required for diet-induced obesity (26), which can further explain the lack of weight gain observed in cisplatin-HFD males, and delayed weight gain in cisplatin-HFD females. Future studies should measure body composition and food consumption of cisplatin- and vehicle-exposed mice to better understand if the observed changes in metabolism correlate with differences in adiposity and caloric intake.

Cisplatin-HFD males displayed low plasma insulin levels through the entirety of the study. Yet, there were no overt differences in insulin content, insulin area, or insulin granularity between islets from cisplatin-HFD and vehicle-HFD males, indicating that the low plasma insulin levels in cisplatin-HFD mice were not driven by depletion of insulin content in β-cells or loss of β-cell mass, aligning with our previous findings in an *in vitro* model (11). It is possible that defects in insulin secretory capacity may be driving low plasma insulin levels in these mice. One group found that HFD-fed β-cell ATP-dependent K^+^ (K_ATP_) channel knockout mice were glucose intolerant, hypoinsulinemic, and more insulin sensitive than controls (27). We previously showed that *in vitro* cisplatin exposure decreased transcription of *Abcc8*, a crucial subunit of the K_ATP_ channel, in mouse islets (11). The combination of cisplatin with HFD-feeding may have exacerbated the effects of cisplatin on K_ATP_ channel subunits, leading to defective nutrient-stimulated insulin release. Understanding the effects of cisplatin on β-cell K_ATP_ channel activity through electrophysiology may help elucidate its impact on insulin secretion.

Interestingly, while the effects of cisplatin on metabolic function *in vivo* were robust and persistent in HFD-fed males relative to chow-fed males, the transcriptional effects of cisplatin on islets were evident in both diet groups. Key β-cell transcriptional regulators (*Pdx1*, *Nkx6-1*, *Mafa*) and functional genes (*Pcsk1/2*, *Ins1*, *Gck*) were downregulated in cisplatin-exposed male islets, irrespective of diet, likely reflecting persistent β-cell injury caused by cisplatin. Despite cisplatin-chow males being metabolically comparable to controls by week 14, the sustained transcriptional changes at week 18 suggest an increased risk of metabolic dysfunction beyond 18 weeks post-exposure. Future studies should determine how cisplatin-exposed mice respond to a metabolic challenge introduced later in life.

Our study focused on phenotyping islet function, but cisplatin is known to cause toxicities to various organs involved in metabolism (28), including the kidney, liver, brain, and GI tract. There were no signs of β-cell loss, α-cell expansion, or increased cytoplasmic proinsulin accumulation in islets from cisplatin-exposed male mice, suggesting that the long-term metabolic effects of cisplatin likely extend beyond the endocrine pancreas. A critical component of glucose homeostasis is the crosstalk between peripheral tissues and pancreatic islets. While we hypothesized that increased hepatic glucose production may be driving hyperglycemia, we observed no robust differences in gene expression of key gluconeogenic and glycogenolytic targets of HFD-fed mice. Comprehensive assessments of glucose uptake, storage, and production in the liver—beyond gene expression—need to be carefully assessed to determine if cisplatin alters hepatic control of glucose homeostasis. Furthermore, previous studies have shown that HFD-feeding disrupts immune function and neuroendocrine signalling (29,30), systems that are also targeted by cisplatin (28). Thus, it is possible that the convergence of these stressors may amplify the adverse effects of cisplatin in HFD-fed mice, leading to prolonged effects of cisplatin on HFD-fed males compared to chow-fed animals.

Similar to our previous findings, we observed minimal sex differences in cisplatin-exposed mice in the 2 weeks post-exposure period (11). However, our current study reveals important sex-specific effects that emerge several weeks after cisplatin exposure. Within the first 4-6 weeks post-exposure, cisplatin led to increased weight loss and impaired compensatory hyperinsulinemia in both male and female HFD-fed mice, as well as elevated fasting blood glucose levels in chow-fed mice of both sexes. Yet, while cisplatin-HFD male mice continued to display a distinctly different metabolic phenotype compared to vehicle-HFD males, cisplatin-exposed female mice recovered and were metabolically similar within diet groups. While most studies showing increased rates of metabolic syndrome following cisplatin treatment focus on male testicular cancer survivors (7,16), one study reported that female ovarian cancer survivors did not have an elevated risk of metabolic syndrome following cisplatin treatment compared to male testicular cancer survivors (31). Despite the limited clinical evidence addressing sex-specific metabolic outcomes of cancer survivors treated with cisplatin, the findings of our mouse study suggest that females may be largely protected from adverse metabolic health outcomes following cisplatin treatment.

Overall, our research indicates that the long-term effects of cisplatin are exacerbated when combined with HFD-feeding and robustly alters the metabolic phenotype of HFD-fed male mice. These findings support a model in which cisplatin disrupts islet adaptation required for diet-induced metabolic compensation, resulting in impaired insulin secretion relative to demand and reduced β-cell plasticity. Overall, our results suggest cancer patients who receive cisplatin and have pre-existing diet-driven metabolic stress may face a higher risk of developing secondary metabolic complications, but their symptoms may not align with a classical T2D phenotype. Importantly, a “Type 5 diabetes” phenotype in cancer survivors—characterized by hyperglycemia, low body weight, low insulin, and lack of insulin resistance—could be easily overlooked or misdiagnosed, highlighting the need to prioritize monitoring metabolic health in cancer survivors following chemotherapy. Moreover, our study highlights the importance of incorporating sex as a biological variable when designing personalized strategies to mitigate long-term metabolic dysregulation in cancer survivors.

## ARTICLE INFORMATION

## Acknowledgements

We sincerely appreciate the support of Carleton Animal Care and Veterinary Services, especially Leslie Peters, Dawson Lafreniere, and Laura Twiner, during this study. We are grateful for Lili Grieco-St-Pierre for thoughtful discussions on data presentation and manuscript revision. We thank Marcia Reid at the Electron Microscopy Facility at McMaster University for her help in sectioning and staining pancreas TEM samples, and Jianqun Wang at the Nano Imaging Facility at Carleton University for his guidance and help in imaging TEM samples. We thank Dr. Melissa Chee at Carleton University for helpful discussions on study design. We appreciate Ineli Perera for her assistance in metabolic assessments, and Nia Tubana-Dean and Emilia Poleo-Giordani for their contributions to image analysis.

## Contribution statement

LB and JEB conceived the experimental design. LB and JEB wrote the manuscript. LB, JP, MEAC, MPH, EvZ, and JEB were involved with data acquisition and analysis. All authors contributed to manuscript revisions and approved the final version of the article.

## Guarantor statement

JEB is the guarantor of this work and, as such, had full access to all the data in the study and takes responsibility for the integrity of the data and the accuracy of the data analysis.

## ABBREVIATIONS

BG: Blood glucose
BW: Body weight
FCCP: Carbonyl cyanide-p-trifluoromethoxyphenylhydrazone
GSIS: Glucose-stimulated insulin secretion
GTT: Glucose tolerance test
HBSS: Hanks’ balanced salt solution
HFD: High-fat diet
HG: High glucose
ITT: Insulin tolerance test
Ip: Intraperitoneal
_KATP_: ATP-dependent potassium
KCl: Potassium chloride
KRBH: Krebs-ringer bicarbonate HEPES buffer
LG: Low glucose
PFA: Paraformaldehyde
PBS: Phosphate buffer saline
T2D: Type 2 diabetes
TEM: Transmission electron microscopy

