## Supplementary material for "Cisplatin exposure alters long-term metabolic phenotype of male, but not female, high-fat diet-fed mice": Suplemental Material

### SUPPLEMENTARY TABLES

**Supplementary Table 1. Immunofluorescence Antibodies Used**

| Antibody | Company (CAT#) | Dilution |
|---|---|---|
| Insulin (C27C9) Monoclonal Antibody | Cell Signalling (#3014) | 1:200 |
| Insulin (L6B10) Monoclonal Antibody | Cell Signalling (#8138S) | 1:250 |
| Glucagon | Sigma-Aldrich (#G2654-0.2mL) | 1:250 |
| Proinsulin | Developmental Studies Hybridoma Bank (#GS-9A8-S) | 1:50 |
| MafA Polyclonal Antibody | Betalogics (#LP9872) | 1:1000 |

**Supplementary Table 2. Primer Sequences for qPCR**

| Target | Species | Forward Sequence (5'-3') | Reverse Sequence (5'-3') |
|---|---|---|---|
| *G6pc* | Mouse | TACTACAGCAACAGCTCCGTG | TCCCAACCACAAGATGACGTT |
| *Gapdh* | Mouse | AGGTCGGTGTGAACGGATTTG | TGTAGACCATGTAGTTGAGGTCA |
| *Gcg* | Mouse | ACTCACAGGGCACATTCACC | CCAGTTTATAAAGTCCCTGG |
| *Gcgr* | Mouse | CTGCACTGCACCCGAAACTA | CACTCAGCCAGATGCTCACA |
| *Gck* | Mouse | TGGTGGATGAGAGCTCAGTG | TGAGCAGCACAAGTCGTACC |
| *Iapp* | Mouse | CAGCTGTCCTCCTCATCCTC | GCACTTCCGTTTGTCCATCT |
| *Ins1* | Mouse | TCAGAGACCATCAGCAAGCA | CTCCCAGAGGGCAAGCAG |
| *Ins2* | Mouse | GCTTCTTCTACACACCCATGT | ACGACTGATCTACAATGCCAC |
| *Mafa* | Mouse | AGTCGTGCCGCTTCAAG | CGCCAACTTCTCGTATTTCTCC |
| *Mafb* | Mouse | TATAAACGCGTCCAGCAGAAGC | CCGGAGTTGGCGAGTTTCTC |

| Target | Species | Forward Sequence (5'-3') | Reverse Sequence (5'-3') |
|---|---|---|---|
| *Nkx6.1* | Mouse | CTTCGCCCTGGAGAAGAC | CCGAGTCCTGCTTCTTCTTG |
| *Pck1* | Mouse | TGGGAACTCACTACTCGGGA | TTCTTCTTGCCTTCGGGGTT |
| *Pcsk1* | Mouse | GGTGGAAAGGTCGAGTCTAGC | TGCACACCAAACGCAAAAGA |
| *Pcsk2* | Mouse | TTTGGAGTCCGAAAGCTCCC | GGTGTAGGCTGCGTCTTCTT |
| *Pdx1* | Mouse | GTACGGGTCCTCTTGTTTTCC | GATGAAATCCACCAAAGCTCAC |
| *Ppia* | Human/Mouse | GCCAGGACCTGTATGCTTTA | AGCTCTGAGCACTGGAGAGA |
| *Pygl* | Mouse | CCAGAGTGCTCTACCCCAAT | CAGCCACCACAAAGTACTCCT |
| *Slc2a2* | Mouse | GCAACTGGGTCTGCAATTTT | CCAGCGAAGAGGAAGAACAC |
| *Slc30a8* | Mouse | GGCTGACATTTGGGTGGTAT | TCACAGGCAAGGTACAGCAG |
| *Sst* | Mouse | CTGAGCAGGACGAGATGAGG | TAACAGGATGTGAATGTCTTCCAGAA |
| *Ucn3* | Mouse | CCCCTCGACCTGAGCATTTC | GGAAGTAGGTGGGCATCAGC |

### SUPPLEMENTARY FIGURES

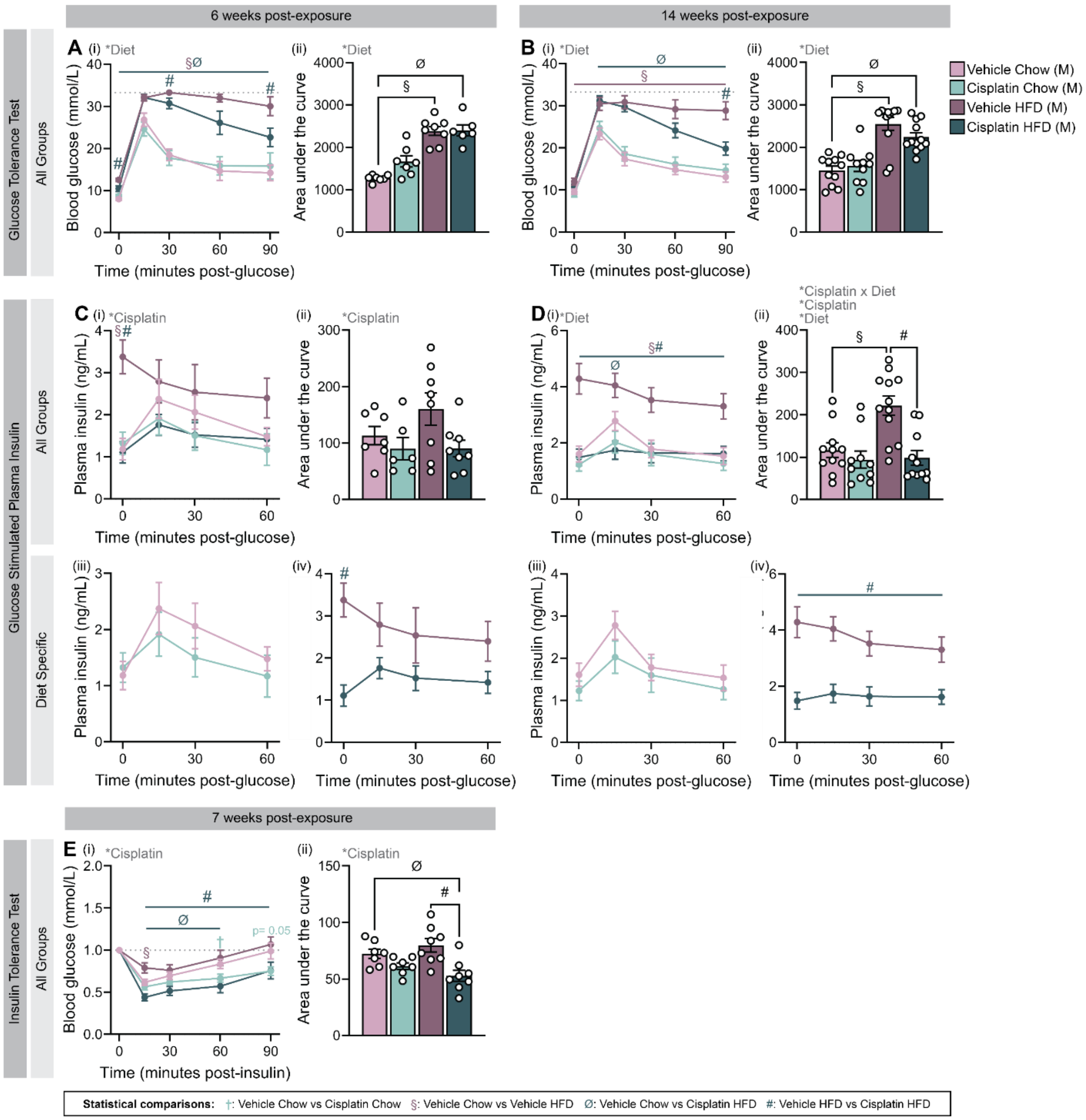


**Supplementary Figure 1. HFD-fed males are glucose intolerant compared to chow-fed males, but cisplatin-HFD mice do not exhibit hyperinsulinemia and are more insulin sensitive compared to vehicle-HFD males. (A-B)** Glucose tolerance and **(C-D)** glucose-stimulated plasma insulin levels were measured at **(A, C)** 6 and **(B, D)** 14 weeks post-exposure. **(E)** Insulin tolerance test conducted at 7 weeks post exposure; blood glucose values are normalized to baseline (t=0). Mice received intraperitoneal injections of **(A, C)** 2 g/kg glucose, **(B, D)** 1 g/kg glucose, or **(E)** 0.6 IU/kg insulin. **(A-B)** Glucose tolerance and **(E)** insulin tolerance are presented as (i) line graphs and (ii) area under the curve. **(C-D)** Plasma insulin levels presented as (i) line graphs with all groups, (ii) area under the curve, (iii) line graphs with only chow-fed groups, and (iv) line graphs with only HFD-fed groups. All data represented as mean ± SEM. Statistical symbols are as follows: *$p < 0.05$ (ANOVA main effects); $^{\dagger}p < 0.05$ (vehicle chow vs. cisplatin chow); $^{\S}p < 0.05$ (vehicle chow vs vehicle HFD); $^{\emptyset}p < 0.05$ (vehicle chow vs. cisplatin HFD); $^{\#}p < 0.05$ (vehicle HFD vs. cisplatin HFD). The following statistical tests were performed: line graphs, repeated measures 3-way ANOVA with Fisher's LSD test; bar graphs, 2-way ANOVA with Tukey's multiple comparisons test. M: Male.

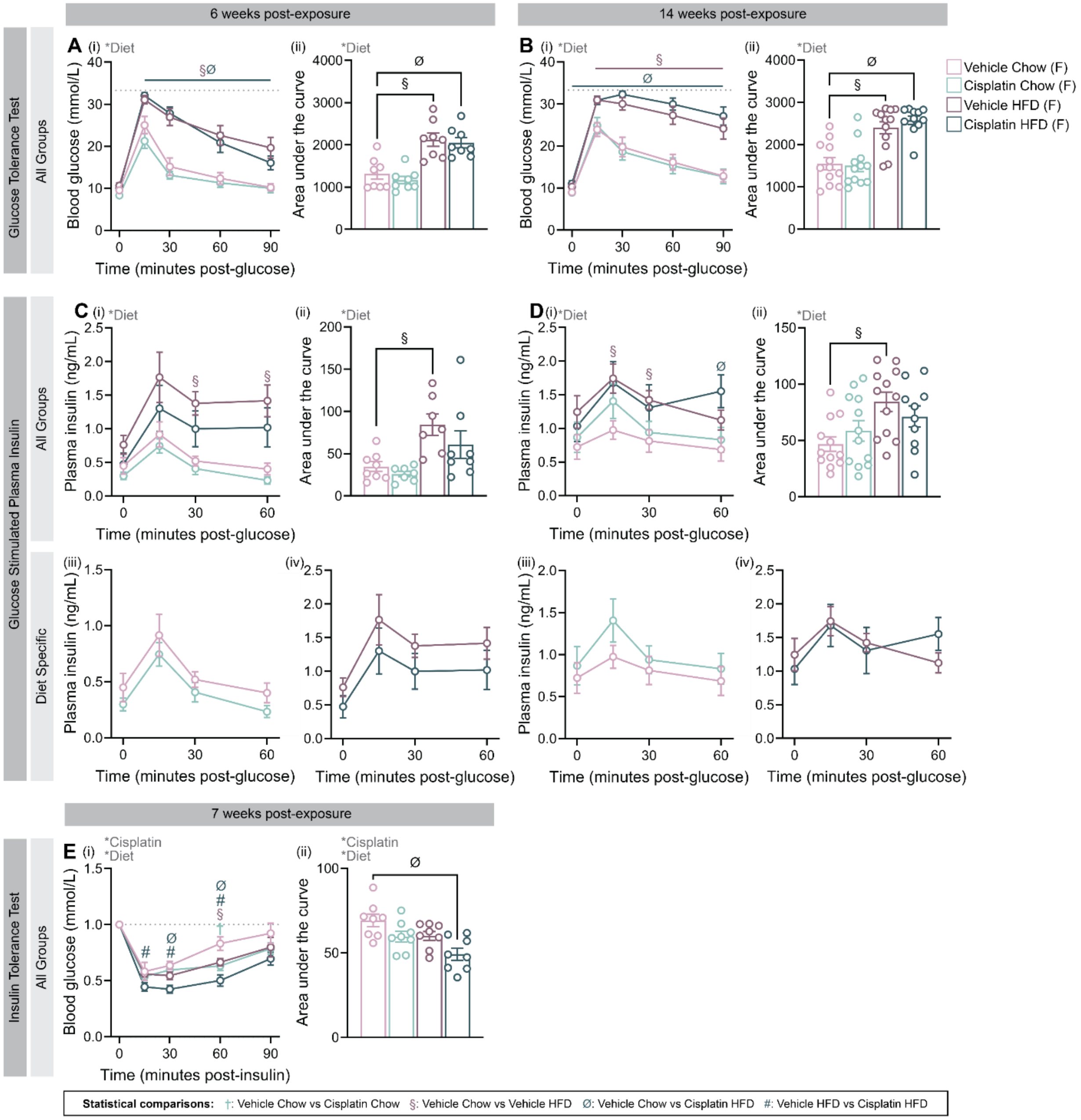


**Supplementary Figure 2. Cisplatin exposure did not alter glucose tolerance or plasma insulin levels in female mice, but increased insulin sensitivity. (A-B)** Glucose tolerance and **(C-D)** glucose-stimulated plasma insulin levels were measured at **(A, C)** 6 and **(B, D)** 14 weeks post-exposure. **(E)** Insulin tolerance test conducted at 7 weeks post exposure; blood glucose values are normalized to baseline (t=0). Mice received intraperitoneal injections of **(A, C)** 2 g/kg glucose, **(B, D)** 1 g/kg glucose, or **(E)** 0.6 IU/kg insulin. **(A-B)** Glucose tolerance and **(E)** insulin tolerance are presented as (i) line graphs and (ii) area under the curve. **(C-D)** Plasma insulin levels presented as (i) line graphs with all groups, (ii) area under the curve, (iii) line graphs with only chow-fed groups, and (iv) line graphs with only HFD-fed groups. All data represented as mean ± SEM. Statistical symbols are as follows: *$p < 0.05$ (ANOVA main effects); $^{†}p < 0.05$ (vehicle chow vs. cisplatin chow); $^{§}p < 0.05$ (vehicle chow vs vehicle HFD); $^{Ø}p < 0.05$ (vehicle chow vs. cisplatin HFD); $^{\#}p < 0.05$ (vehicle HFD vs. cisplatin HFD). The following statistical tests were used: line graphs (A-C, E) repeated measures 3-way ANOVA with Fisher's LSD test, (D) repeated measures mixed-effects analysis with Fisher's LSD test; bar graphs, 2-way ANOVA with Tukey's multiple comparisons test. F: Female.

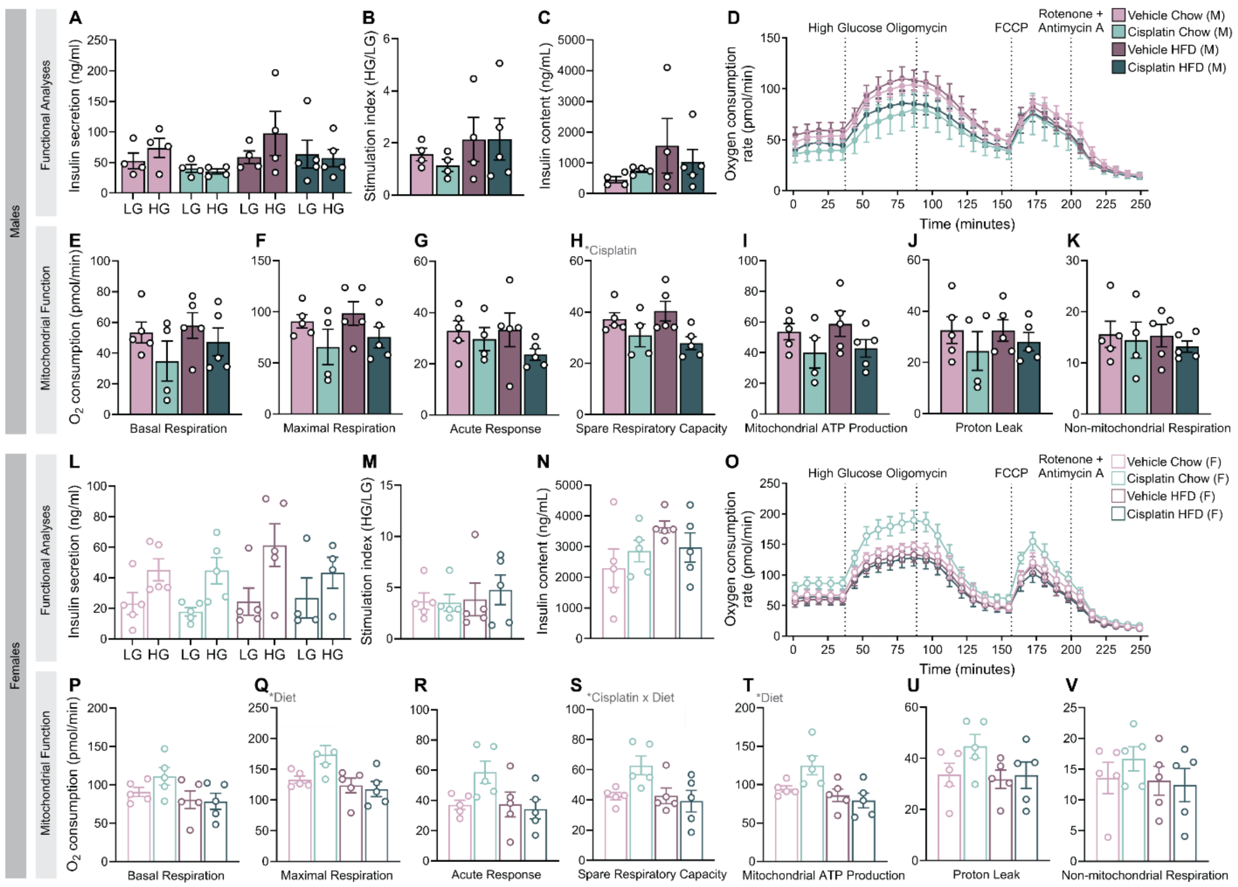


**Supplementary Figure 3. Cisplatin exposure did not alter glucose-stimulated insulin secretion or oxygen consumption in islets isolated 18 weeks post-exposure.** Islets were isolated from **(A-K)** male and **(L-V)** female mice 18 weeks post-exposure (see Figure 1A). **(A, L)** Static glucose-stimulated insulin secretion was assessed via sequential 1-hour incubations in low glucose (LG; 2.8 mmol/L) and high glucose (HG; 16.7 mmol/L) buffer. **(B, M)** The stimulation index was calculated as a ratio of insulin secretion under HG conditions to LG conditions. **(C, N)** Islets were lysed in acid ethanol and total insulin content was measured. **(D, O)** Oxygen consumption was measured using a Seahorse XFe24 analyzer. Islets were incubated in LG media then exposed to serial injections of 16.7 mmol/L glucose, 2.5 µmol/L oligomycin, 3 µmol/L carbonyl cyanide-p-trifluoromethoxyphenylhydrazone (FCCP), and a combination of 3 µmol/L rotenone and antimycin A. **(E-K, P-V)** Parameters of mitochondrial function including **(E, P)** basal respiration, **(F, Q)** maximal respiration, **(G, R)** acute glucose response, **(H, S)** spare respiratory capacity, **(I, T)** mitochondrial ATP production, **(J, U)** proton leak, and **(K, V)** non-mitochondrial respiration. All data represented as mean ± SEM. Statistical symbols are as follows: *$p < 0.05$ (ANOVA main effects); $^{\dagger}p < 0.05$ (vehicle chow vs. cisplatin chow); $^{\S}p < 0.05$ (vehicle chow vs vehicle HFD); $^{\emptyset}p < 0.05$ (vehicle chow vs. cisplatin HFD); $^{\#}p < 0.05$ (vehicle HFD vs. cisplatin HFD). The following statistical analyses were done: (A, D, L, O) repeated measures 3-way ANOVA with Fisher's LSD test, (B-C, E-K, M-N, P-V) 2-way ANOVA with Tukey's multiple comparisons test. M: Male, F: Female.

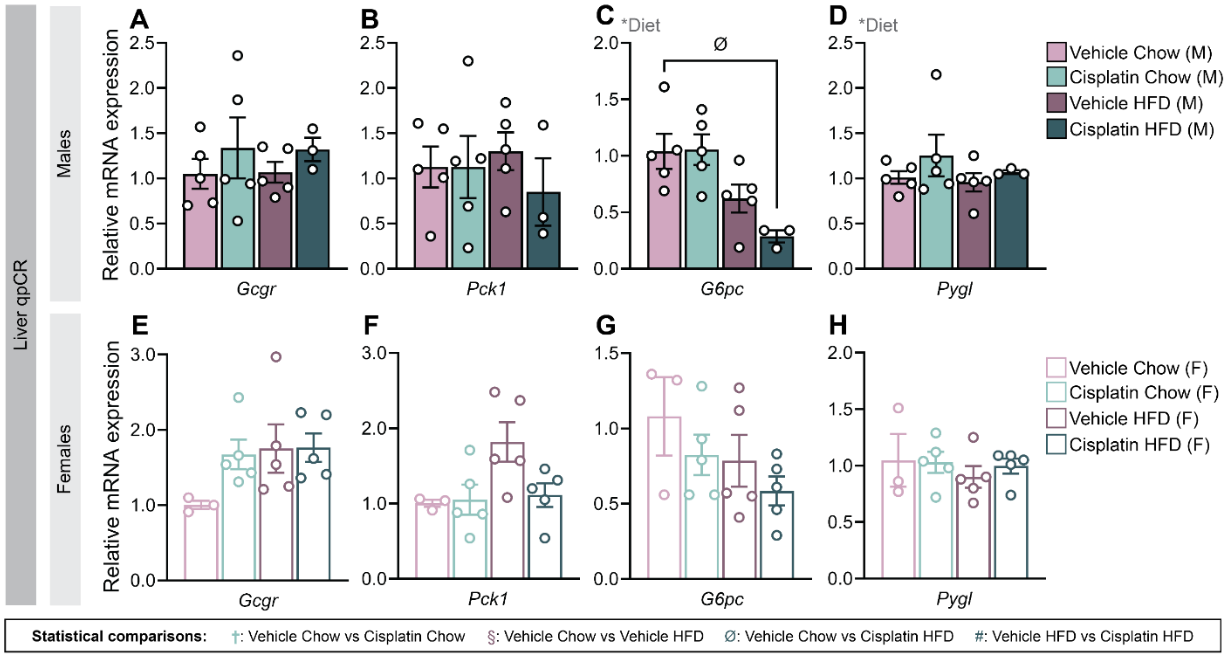


**Supplementary Figure 4. Expression of key genes in the liver linked to metabolic function were largely unaffected by cisplatin or diet in mice.** Flash frozen liver from **(A-D)** male and **(E-H)** female mice 18 weeks post exposure were used to measure relative mRNA expression of **(A, E)** *Gcgr*, **(B, F)** *Pck1*, **(C, G)** *G6pc*, and **(D, H)** *Pygl*. All data represented as mean ± SEM. Statistical symbols are as follows: $*p < 0.05$ (ANOVA main effects); $^{\dagger}p < 0.05$ (vehicle chow vs. cisplatin chow); $^{\S}p < 0.05$ (vehicle chow vs vehicle HFD); $^{\varnothing}p < 0.05$ (vehicle chow vs. cisplatin HFD); $^{\#}p < 0.05$ (vehicle HFD vs. cisplatin HFD). All graphs analyzed using 2-way ANOVA with Tukey's multiple comparisons test. M: Male, F: Female.
